# Maternal behavioural thermoregulation facilitated evolutionary transitions from egg laying to live birth

**DOI:** 10.1101/2021.02.07.430163

**Authors:** Amanda K. Pettersen, Nathalie Feiner, Daniel W.A. Noble, Geoffrey M. While, Tobias Uller, Charlie K. Cornwallis

**Affiliations:** Department of Biology, Lund University, Sweden; School of Life and Environmental Sciences, The University of Sydney, Australia; Division of Ecology and Evolution, Research School of Biology, The Australian National University, Australia; School of Natural Sciences, University of Tasmania, Australia

**Author notes:** Joint senior author (Tobias Uller, Charlie K. Cornwallis).

## Abstract

Live birth is a key innovation that has evolved from egg laying ancestors over 100 times in reptiles. However, egg-laying lizards and snakes often possess preferred body temperatures that are lethal to developing embryos, which should select against egg retention. Here, we demonstrate that thermal mismatches between mothers and offspring are widespread across the squamate phylogeny. This mismatch is resolved by gravid females adjusting their body temperature towards the thermal optimum of embryos. Importantly, phylogenetic reconstructions suggest this thermoregulatory behaviour evolved in egg-laying species prior to the evolution of live birth. Maternal thermoregulatory behaviour therefore bypasses the constraints imposed by a slowly evolving thermal physiology and has likely been a key facilitator in the repeated transitions to live birth.

## Introduction

The evolution of live birth is an important life-history adaptation in vertebrates^1–3^. The ecological conditions that favour the transition from egg laying (oviparity) to live birth (viviparity) are relatively well understood, especially in reptiles, with particularly strong support for the adaptive value of viviparity in cool climates^3–5^. By retaining embryos throughout development, mothers can buffer offspring from suboptimal nest temperatures, ensuring faster development, higher hatching success, and increased offspring viability^4,6–8^. This transition has allowed reptile species to diversify and persist in cool climates across the globe^9^.

Despite clear adaptive advantages, the evolutionary transition from oviparous ancestors to viviparity is challenging to explain. Evidence from case studies of lizards and snakes show that embryos and adults of oviparous species have different thermal requirements, with adult preferred body temperatures often exceeding the upper lethal limit of embryos^6,10–12^. For example, the average nest temperature of the Iberian emerald lizard, *Lacerta schreiberi*, is 24°C, rarely exceeding 30°C, whereas the preferred body temperature of females is 33°C^13^. Since embryos are well adapted to the temperatures they typically experience in the nest, they generally have limited capacity to develop at temperatures outside of this range^14–16^. Prolonged exposure to temperatures optimal for adult female performance should therefore result in offspring abnormality and death^15,17–19^.

A mismatch between thermal optima of embryos and adult females should prevent mothers from retaining embryos throughout development, inhibiting evolutionary transitions to live birth^20^. Despite this apparent constraint, live birth has evolved over 100 times in squamate reptiles^21,22^. How can we reconcile the repeated evolution of viviparity with the potentially widespread thermal mismatches between embryos and adults in oviparous species? One possibility is that females behaviourally adjust their body temperature when pregnant to close the gap between adult and embryo thermal optima, even when substantial mismatches in thermal preferences exist. Such plasticity may temporarily come at a cost to female performance but shifting body temperatures while gravid to match the thermal optima of embryos could eliminate thermal barriers to the evolution of viviparity. An alternative possibility is that viviparity only evolves from oviparity in species where there is no mismatch, that is, where adult body temperature and embryo thermal optima are well aligned.

Here we show that thermal mismatches between mothers and offspring are widespread across the squamate phylogeny. We use phylogenetic comparative analyses to examine if maternal thermoregulatory behaviour has eliminated thermal mismatches between embryos and adults, enabling the repeated evolution of viviparity across reptiles. First, we test whether adult females adjust their body temperature when gravid to better match the temperature optimum of their developing embryos. To this end, published data was collated on the thermal preferences of non-gravid and gravid females and the thermal tolerance of embryos. Using meta-analytical techniques, we calculated a standardised effect size (Hedges’ *g*) of the amount females change their preferred body temperature when gravid (Tables S1 & S2). Second, we test whether behavioural plasticity was more pronounced in viviparous species compared to oviparous females, as expected if thermal conflicts are more severe, and therefore avoided by species that are egg-laying. Third, we test the alternative possibility that viviparity evolves in lineages where adult and embryo optima are aligned using ancestral reconstructions.

## Results

### Female behavioural plasticity resolves the constraints imposed by thermal physiology

Across 52 species of reptiles (N_viviparous species_= 32, N_oviparous species_= 20) mismatches between the preferred temperatures of non-gravid females (*P*_*bt-ng*_) and embryos (*T*_*opt*_) were widespread. On average *P*_*bt-ng*_ of adult females was 4 °C higher than the embryonic *T*_*opt*_. We found that females significantly altered their body temperature when gravid (*P*_*bt-g*_) to reduce such mismatches.

Specifically, in species with high non-gravid preferred body temperature (*P*_*bt-ng*_), where thermal conflicts are potentially most severe, females significantly reduced their body temperature when gravid (negative values of Hedges’ *g*; Fig. 1). Conversely, in species with low preferred body temperatures females increased their body temperature when gravid (positive values of Hedges’ *g*; Fig. 1). Combined this strongly suggests that females with extreme body temperatures regulate their own body temperature to meet the thermal optima of embryos (Table S3 = M1). This appears to be required as there was little evidence that the optimal temperature for embryo development (*T*_*opt*_) coevolves with non-gravid female preferred body temperatures (phylogenetic correlation (MR-BPMM): PM = 0.53, CI: -0.27, 0.93, *pMCMC* = 0.15. Table S4 = M2) and both *P*_*bt*_ and *T*_*opt*_ were estimated to evolve slowly (raw data mean±SD: Oviparous species, *P*_*bt*_ = 32.41±4.15 °C, N_species_ = 103; *T*_*opt*_ = 27.15±1.92 °C, N_species_ = 47. Viviparous species *P*_*bt*_ = 29.5±4.06 °C, N_species_ = 61; *T*_*opt*_ = 26.0 ±2.23 °C, N_species_ = 5. MR-BPMM: *P*_*bt*_ phylo *H*^2^: 0.91, CI: 0.80, 0.95. *T*_*opt*_ phylo *H*^2^: 0.91, CI: 0.71, 0.99. Table S4 = M2).

**Fig. 1.**
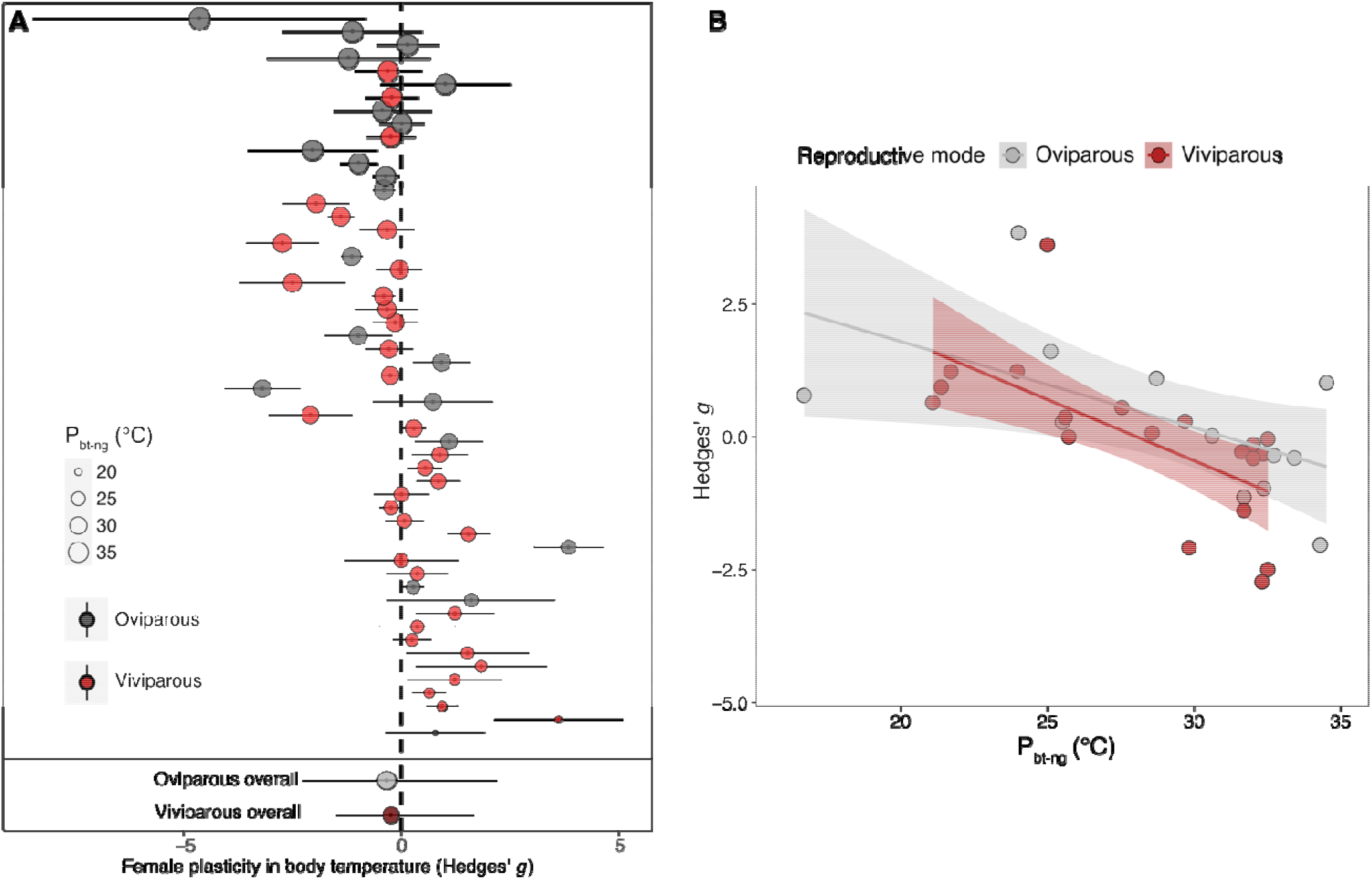
Plasticity in female body temperatures when gravid (Hedges’ *g*) resolves thermal mismatches between adults and embryos across 52 extant oviparous (n=20) and viviparous (n=32) squamate reptiles. (**A**) Adjustment of body temperature in gravid females (*P*_*bt-g*_) in relation to their non-gravid preferred body temperature (*P*_*bt-ng*_); Hedges’ *g*; *P*_*bt-g*_ - *P*_*bt-ng*_. Data points are ordered along the y axis according to *P*_*bt*-ng_. Points represent species means ± SEs and the size of points is scaled to indicate *P*_*bt*-ng_ (°C). (**B**) Relationship between Hedges’ *g* and *P*_*bt-ng*_. Regression lines ± 95% credible intervals are plotted. High values of Hedges’ *g* signify an increase in body temperature when gravid while low values imply a decrease. Positive values of Hedges’ *g* indicate higher gravid (*P*_*bt-g*_) versus non-gravid (*P*_*bt-ng*_) and negative values indicate reduced *P*_*bt*_ when gravid (*P*_*bt-g*_) compared with non-gravid (*P*_*bt-ng*_). The plots shows that species with high *P*_*bt-ng*_ tend to reduce their body temperature when gravid (negative Hedges’ *g*), whereas species with low *P*_*bt-ng*_ tend to increase their body temperature when gravid (positive Hedges’ *g*).

### Gravid females shift their body temperatures towards embryo thermal optima regardless of parity mode

Contrary to the expectation that selection for behavioural plasticity is greater in live-bearing females, their adjustment of body temperature when gravid did not differ from egg-laying females (Table S5 = M3). Egg-laying females with higher *P*_*bt*_ down-regulated their temperature when gravid while species with low *P*_*bt*_ up-regulated their body temperatures in a similar way to live-bearing females (Fig. 2. Phylogenetic correlation between *P*_*bt*_ and Hedges’ *g*: Oviparous PM = -0.90, CI: -0.97, 0.02, *pMCMC* = 0.05; Viviparous PM = -0.88, CI: -0.98, -0.30, *pMCMC* = 0.01. Table S6 = M4). Ancestral reconstructions of Hedges’ *g* also showed that, in lineages where there were thermal mismatches between adults and embryos, females adjusted their body temperature to a much greater extent than when their thermal optima were aligned, irrespective of whether they were egg-laying or live-bearing (Fig. 2. MR-BPMM: Oviparous ancestors Hedges’ *g* PM = -1.22, CI = -3.09, -0.44, *pMCMC* < 0.01. Viviparous ancestors Hedges’ *g* PM = -1.27, CI = -1.63, -0.53, *pMCMC* = 0.001. Table S7 = M5). Consequently, estimates of Hedges’ *g* did not differ between the ancestors of oviparous and viviparous species (Fig. 2B). This suggests that behavioural plasticity was present prior to the emergence of live birth.

**Fig. 2.**
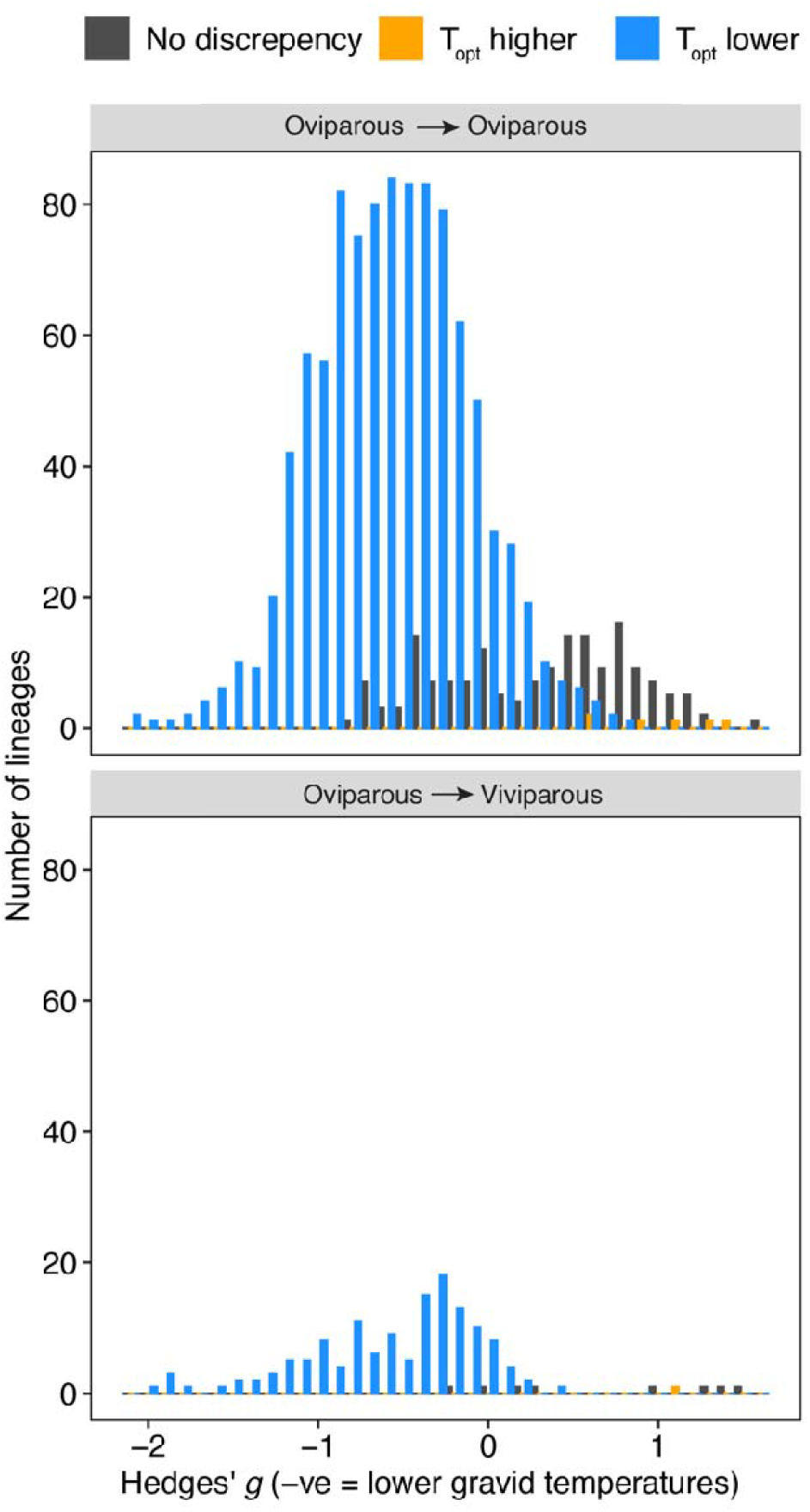
Female behavioural plasticity facilitates transitions to live birth. The adjustment of body temperature by females in lineages where egg laying was maintained (top panel) and where live bearing evolved (bottom panel) in relation to whether embryos had a significantly lower (T_opt_ lower; blue), higher (T_opt_ higher; orange), or aligned (No discrepancy; grey) estimated thermal optimum than adult preferred body temperature (see Methods section “*Testing if Hedges’ g is related to discrepancies between P*_*bt*_ *and T*_*opt*_ *in the ancestors of oviparous and viviparous species*” for how mismatches in thermal optima were estimated).

### An alternative explanation for the evolution of viviparity from oviparity?

The presence of female thermal plasticity in egg-laying and live bearing species suggests it may circumvent the barriers to the evolution of live-bearing imposed by mismatches in slowly evolving thermal optima of adults and embryos. However, another possibility is that viviparity evolves predominantly in lineages where adult and embryo thermal optima are already aligned, alleviating the potential costs to females of adjusting their body temperatures. Estimates of *P*_*bt*_ and *T*_*opt*_ in the egg-laying ancestors of live-bearing species showed that they were no more likely to have aligned adult and embryo thermal optima than the ancestors of egg-laying species. Specifically, 5% of ancestors of live-bearing species had aligned embryo and adult thermal optima compared to 14% in the ancestors of egg-laying species (Fig. 3, χ^2^ = 3.6, df=1, P > 0.05. Table S8-S9 = M5). Consequently, in 95% of the oviparous ancestors of viviparous species there were mismatches between the predicted thermal optima of embryos and adults, illustrating a widespread need for female plasticity to resolve thermal conflicts (Table S8-S9).

**Fig. 3.**
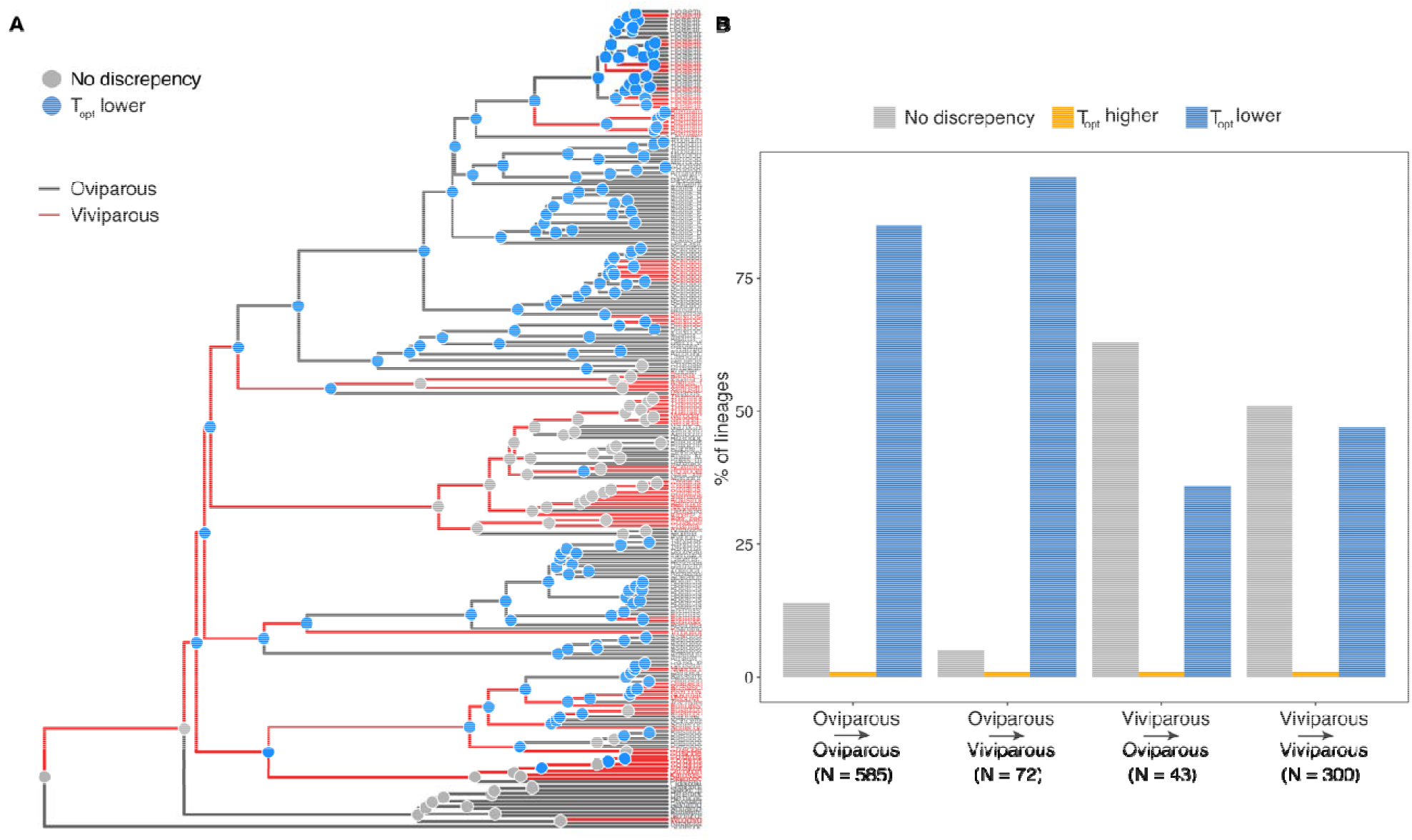
Alignment of embryo and adult thermal optima and transitions to viviparity across 224 species of squamate reptiles. (**A**) Tip labels and branches are coloured according to reproductive modes (red = live bearing/viviparous, grey = egg laying/oviparous; branch colours represent predicted ancestral values; Table S9). Coloured nodes correspond to the discrepancy between *P*_*bt*_ and *T*_*opt*_ (grey = no discrepancy, blue = *T*_*opt*_ < *P*_*bt*_). (**B**) Percentage of lineages with transitions between reproductive modes and discrepancy between *P*_*bt*_ and *T*_*opt*_ (grey = no discrepancy, blue = *T*_*opt*_ < *P*_*bt*_, orange = *T*_*opt*_ > *P*_*bt*_).

## Discussion

Our results suggest that the upper thermal limit of embryos is commonly lower than the preferred body temperature of females in oviparous snakes and lizards. Both adult and embryo thermal biology appear to evolve slowly, generating a wide-spread and evolutionarily persistent mismatch between the thermal optima of mothers and their embryos. Our data suggest non-gravid females have preferred body temperatures that are on average 4 °C higher than the temperature that maximises hatching success. Incubation experiments have shown that exposure to such temperatures throughout development may cause malformations or jeopardise embryo survival (reviewed in^15^). As there was little evidence that female preferred body temperature and offspring thermal optima coevolve, such mismatches may be difficult to resolve and thus hamper transitions to viviparity.

Our findings support the hypothesis that the thermal mismatch between females and embryos is resolved by females adjusting their body temperature when gravid to meet the thermal requirements of their embryos^23^. This behavioural plasticity effectively eliminates barriers to the evolution of viviparity. The shifts in body temperature between gravid and non-gravid females (Hedges’ *g*) were significantly phylogenetically correlated with the discrepancy between adult and embryo thermal optima in both oviparous and viviparous species. Moreover, ancestral state reconstructions suggest that this behavioural plasticity was present prior to the emergence of live birth, negating the need for adult and embryo thermal optima to be aligned for viviparity to evolve.

The down-regulation of body temperature by gravid egg-laying females may appear surprising considering that most of these species lay their eggs within the early stages of development (commonly around the time of limb bud formation)^24^. However, early developmental stages, involving gastrulation, neurulation and organogenesis, are potentially even more sensitive to thermal stress than later stages, which are predominantly associated with growth^16,20^. The temperature sensitivity of early-stage embryos may therefore generate selection for the resolution of mother-offspring thermal conflicts in both egg-laying and live-bearing species. If true, the key innovation of live birth may owe its evolutionary origin to mechanisms of behavioural temperature regulation put in place long before live birth emerged.

Behavioural plasticity has continued to play an important role in thermal adaptation over and above facilitating the evolution of live birth. Specifically, behavioural plasticity is frequently maintained in viviparous species that have colonised cool climates. Such behavioural flexibility enables females to upregulate their body temperature to maintain embryos at significantly warmer temperatures than the external environment, contributing to the adaptive value of viviparity ^2,7,8,25,26^. In turn, the ability to cope with a greater range of temperature conditions has the potential to allow populations to persist and expand into suboptimal environments^27^. Female thermoregulatory behaviour therefore appears to be a key adaptation that helps resolve thermal mismatches between adults and embryos and facilitate the expansion of reptiles into a variety of environments.

## Acknowledgments

This work was funded by a Wenner-Gren Foundation Postdoctoral Fellowship (UPD2019-0208) to AKP and TU, a Wallenberg Academy fellowship (2018.0138) to CKC and starting grants from the European Research Council (#948126) and the Swedish Research Council (#2020-03650) to NF.

## Competing interests

Authors declare that they have no competing interests.

### Data and materials availability

All data and code are publicly available at: https://doi.org/10.17605/OSF.IO/JT28V

### Methods

#### Literature search and data collection

To investigate the relationship between maternal behavioural plasticity and embryo thermal sensitivity, we used reproductive mode data from Pyron and Burbrink^28^, and collated existing data to collate three datasets: 1) female body temperature when gravid (*P*_*bt-g*_) and not gravid *(P*_*bt- ng*_) (“Hedges’ *g* dataset”); 2) female preferred body temperature (*P*_*bt*_) (“*P*_*bt*_ dataset”); and 3) the temperature at which hatching success was maximised (*T*_*opt*_) (“*T*_*opt*_ dataset”). Complete data are presented in Table S1. Data were collated for each of the variables from literature searches using ISI *Web of Science* (v.5.30) with search terms specific to each dataset. Search results were then imported and sorted for relevance using Rayyan software^29^ (for details see Supplementary Information).

#### Estimating Hedges’ g

Published articles presenting data on thermal preference (preferred body temperature) in gravid (*P*_*bt-g*_) versus non-gravid (*P*_*bt*-ng_) adult female squamate species were collected for the “Hedges’ *g*” dataset, following the preferred reporting items for systematic reviews and meta-analyses (PRISMA) statement^30^. We conducted a literature search using ISI *Web of Science* (v.5.30) with the ‘title’, ‘abstract’ or ‘keywords’ search terms ‘body temperature*AND gravid*OR reproduct*’, along with one of the following: ‘squamat*’, ‘lizard*’, ‘snake*’ which yielded a total of 721 papers. We searched available literature for both field and laboratory measures of female body temperature, comprising studies that directly compared *Pbt*_*gravid*_ and *Pbt*. 648 papers were rejected due to irrelevance (PRISMA statement; Fig. S1). We only included studies that provided both sample size and error around mean preferred body temperature. Studies used artificial temperature gradients in the laboratory (n = 37) or measured preferred basking temperature in the field (n = 36). Laboratory studies generally measured body temperature in the same female during gestation (*P*_*bt-g*_) and either before or after gestation (*P*_*bt-ng*_) as repeated measures. In contrast, field studies often measured body temperature on a population during the reproductive season, comparing body temperatures in gravid and non-gravid females at a single time point. Combined, this yielded a total of 73 studies published up to July 2022 from which effect sizes were calculated for 52 species (live bearing: n = 32 and egg laying: n = 20). Effect sizes of female adjustment of body temperature when gravid for each species were calculated as the standardised mean differences (Hedges’ *g*) in preferred body temperatures between non-gravid and gravid females (*P*_*bt-g*_ - *P*_*bt-ng*_), adjusting for small sample sizes^31^. We examined the mean Hedges’ *g* between laboratory or field studies and found there were no significant differences (PM (Oviparous) = -0.67, CI = -1.63, 0.60; PM (Viviparous) = 0.65, CI = -0.20, 1.30. Table S10. See Verification analyses in Supplementary Information).

#### Estimating P_bt_ independently from Hedges’ g

We collected independent data on the preferred body temperature in adult females (*P*_*bt*_) using the same method described for the “Hedges’ *g*” dataset, using search terms ‘body temperature*’, along with one of the following: ‘squamat*’, ‘lizard*’, ‘snake*’ which yielded a total of 1075 papers. We only used data from studies where *Pbt* for females was stated explicitly (unless pooled male/female data stated no significant effect of sex) and excluded data on females that were described to be gravid or data collected during the reproductive season. We additionally cross-referenced this search with articles cited in^32^, supplementing our original dataset with 42 studies (PRISMA statement; Fig. S2). This provided a final dataset of female preferred body temperature for 163 species that was independent of the gravid and non-gravid measures used to calculate Hedges’ *g* (live bearing: n = 61 and egg laying: n = 103, note *P*_*bt*_ data for *Zootoca vivipara* was available for both reproductive modes and both were included in the analyses – see below).

#### Estimating Topt

Hatching success and egg incubation temperature data for 47 egg-laying and 4 live-bearing species was extracted from the Reptile Development Database (RepDevo vers 1.0.2; ^33^), and from the literature using search terms: ‘temperature*AND incubat*AND hatch*OR surv*’, along with one of the following: ‘squamat*’, ‘lizard*’, ‘snake*’, yielding a total of 671 papers (PRISMA statement; Fig. S3). We only included studies where three or more constant temperature treatments were used under controlled laboratory conditions, resulting in 661 papers being rejected due to irrelevance or overlap with the Reptile Development Database. The final *Topt* dataset, consisting of 51 species, obtained from 81 studies. Thermal performance curves, relating hatching success with incubation temperature, were fit using nonlinear least-squares regression with the *nls*.*LM* function in the *minpack*.*LM* package in *R*^34^. For each species-specific function we then calculated a single-point estimate at which optimal hatching success occurred (here-on designated “Topt”). Raw thermal performance data are provided in Fig. S4.

Given that *T*_*opt*_ had a strong phylogenetic signature (Phylogenetic Heritability (*H*^*2*^) = 0.95, 95% CI: 0.74 – 0.99; Table S4) we fitted a Bayesian Phylogenetic Mixed Effects Model (BPMM) to hatching success data for all species jointly. Including phylogenetic information allowed for the *T*_*opt*_ of each species to be estimated with greater accuracy and precision given that the range and number of temperatures across species varied (range: 10-40 °C, mean number of temperatures per species ± SD: 7.62 ± 4.29). Compared to non-phylogenetic models, *T*_*opt*_ estimates produced from phylogenetic models (BPMM) showed smaller sampling error and avoided convergence problems in estimating model parameters.

Importantly, this approach was not used to estimate *T*_*opt*_ data for species without data, only to better predict *T*_*opt*_ values for species for which there were data. The *T*_*opt*_ BPMM model was run for 1,100,000 iterations with a burn-in of 100,000 iterations and thinning rate of 500, leaving us with 2,000 samples from the posterior distribution. Autocorrelation was low (lag values < 0.1) and trace plots showed chains mixed well for all parameters. Our model included temperature as a fixed effect (estimating both a linear and quadratic slopes) and random slopes of temperature (linear and quadratic slopes) fitted at the phylogenetic level. From our BPMM we estimated *T*_*opt*_, and its corresponding sampling variance, using the posterior distribution of fixed effects and best linear unbiased predictors (BLUPs) for the random slopes (linear and quadratic) for each species as follows:

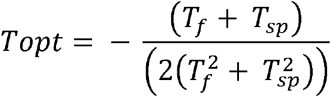

Where *T*_f_ and 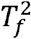 are the posterior linear and quadratic fixed effect estimates for temperature and *T*_sp_ and 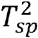 are the posterior BLUPs for a given species extracted from the phylogenetic random slopes. Calculating *T*_*opt*_ using the posterior distribution of fixed and random effects meant that sampling error for a given species could be propagated to subsequent analyses (see below).

## General Statistical Methods

We used Bayesian Phylogenetic Mixed Effects Models with single (BPMM) and multiple response variables (MR-BPMM) to estimate phylogenetic correlations between traits and reconstruct ancestral states of continuous variables. In all MR-BPMM global intercepts were removed to estimate an intercept for each trait. Hidden Markov Models (HMMs) were used to reconstruct ancestral states of viviparity and Phylogenetic Ridge Regression (PRR) were used to check for rate shifts in continuous traits across the phylogenetic tree. All analyses were conducted in R version 4.0.1^35^.

### Bayesian Phylogenetic Mixed Effects Models (BPMM)

We implemented BPMMs in *R* with the *MCMCglmm* package^36^. Hedges’ *g, P*_*bt*_ *and T*_*opt*_ were modelled with Gaussian error distributions. For some species there were multiple estimates of Hedges’ *g*, which was accounted for in two ways. In multi-response models where relationships between multiple traits were examined, a single data point of a weighted mean of Hedges’ *g* was included for each species. In models where Hedges’ *g* was a single response variable (M3 and M6), species was included as a random effect to account for multiple data points per species.

The random effect *animal* with a variance-(co)variance matrix derived from the phylogenetic tree was included in all models^36^. We calculated the phylogenetic signature (equivalent to heritability, *H*^2^, in the terminology of *MCMCglmm*) for each trait as the variance explained by *animal* relative to total random effect variance. Multi-response BPMMs (MR-BPMMs) fitted with *MCMCglmm* allow the phylogenetic and within-species (residual) correlations between traits to be estimated by fitting unstructured covariance matrices. Correlations between traits (e.g., A & B) were calculated as:

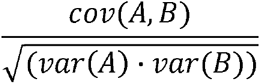

### Model convergence, prior settings and characterisation of posterior distributions

Non-informative uniform priors were used for fixed effects and inverse-Wishart priors for random effects (*V* = 1, nu = 0.002;^36^). To examine model convergence we ran three independent MCMC chains and examined autocorrelation, which was low (lag values < 0.1), trace plots, which showed chains mixed well, and Gelman and Rubin’s convergence diagnostic that models converged (potential scale reduction factors were all below 1.1: R function gelman.diag^37^). All models were the run for 3000000 iterations, with a burn-in of 999500 iterations and every 2000^th^ iteration was saved for parameter estimation (see accounting for phylogenetic uncertainty and data imputation section for more details). Posterior distributions of all parameters were characterised using modes and 95% credible intervals (CIs). Effects were regarded as significant where CIs did not span 0. *pMCMC* (number of iterations above or below 0 / total number of iterations) are also presented to facilitate general interpretation.

### Missing data across species

BPMMs permit missing data in response variables which was crucial given the patchy distribution of the data (Table S1). The accuracy with which missing data is predicted is related to the phylogenetic signature in traits and the strength of phylogenetic correlations between traits^38^. All traits had high phylogenetic signature (phylogenetic H^2^ >70%) producing high correspondence between raw and predicted values (Fig. S6; See also Supplementary Information). As a result, our BPMMs enabled us to deal with the fact that not all traits have been measured in all species.

### Accounting for differences in sampling variances across data points

The accuracy of measures of Hedges’ *g, P*_*bt*_ and *T*_*opt*_ varied across species due to study design and sample sizes which can be accounted for by weighting data points by their inverse sampling variance using the ‘mev’ term in *MCMCglmm*. However, missing values in sampling variances are not permitted in *MCMCglmm*. As data on the error and sample size was missing for Hedges’ *g, P*_*bt*_ and *T*_*opt*_ it would not have been possible to account for sampling error in our analyses without drastically reducing the size of our dataset. Consequently, we used multiple imputation with predictive mean matching in the *mice* package in R to impute missing error and sample sizes^39^. Samples sizes were not available for reproductive mode, but it is typically invariant within populations leading to minimal measurement error. Therefore, the mev term for reproductive mode was specified as 0.

To incorporate uncertainty in imputations, 20 complete datasets were generated, and all analyses were conducted by sampling across these datasets. Each model sampled through the 20 datasets 75 times (1500 sampling events) for 2000 iterations with only the last iteration being saved. Estimates from the last iteration of each sampling event *i* were used as the starting parameter values for the next *i* + 1. This led to a posterior sample of 1500 iterations, the first 500 iterations were discarded as a burn-in and the remaining 1000 (50 per dataset) were used to estimate parameters (total iterations = 3000000 (2000 x 1500), burn-in = 999500 (1999 x 500)). Pooling of posterior distributions from model parameters from across each of the *n = 20* datasets enabled imputation uncertainty in sampling variances to be accounted for in the posterior distribution.

### Phylogeny and accounting for phylogenetic uncertainty

We used a recent phylogeny of squamates pruned to the 224 species with thermal data^40^, For BPMMs each model sampled through 1500 trees using the same procedure as described for sampling across imputed datasets. Each of the 1500 posterior samples was obtained using a different tree. Pooling of posterior distributions from model parameters from across trees enabled phylogenetic uncertainty to be accounted for. To account for phylogenetic uncertainty in the reconstructions of viviparity, HMMs were run on the same 1000 trees that posterior estimates were obtained from using BPMMs (see ‘*Testing if Hedges’ g is related to discrepancies between P*_*bt*_ and *T*_*opt*_ *in the ancestors of oviparous and viviparous species’* for more details). For figures and to compare the performance of different analytical techniques in reconstructing reproductive mode, *T*_*opt*_ and *P*_*bt*_ we used the maximum clade credibility tree provided by^40^.

## Specific Statistical Analyses

### Testing if Hedges’ g is related to P_bt_

The phylogenetic correlation between *P*_*bt*_ and Hedges’ *g* was estimated using a MR-BPMM with unstructured phylogenetic and residual covariance matrices fitted as random effects (Table S3. R code model M1.1).

### Estimating the phylogenetic correlation between P_bt_ and Topt

The phylogenetic signature in *P*_*bt*_ and *T*_*opt*_ and their phylogenetic correlation was estimated using a MR-BPMM with unstructured phylogenetic and residual covariance matrices fitted as random effects (Table S4. R code model M2.1).

### Testing if Hedges’ g is different between oviparous and viviparous species

Differences in Hedges’ *g* between oviparous and viviparous species were tested using a BPMM with reproductive mode as a fixed effect (Table S5. R code model M3.1).

### Testing if the relationship between Hedges’ g and P_bt_ differs between oviparous and viviparous species

We tested if the relationship between Hedges’ *g* and *P*_*bt*_ differed between oviparous and viviparous species using a MR-BPMM with separate unstructured phylogenetic and residual covariance matrices for each reproductive mode specified using the ‘at.level’ function in *MCMCglmm* (Table S6. R code model M4.1).

### Testing if Hedges’ g is related to discrepancies between P_bt_ and Topt in the ancestors of oviparous and viviparous species

To examine values of Hedges’ *g* in relation to the discrepancy between *P*_*bt*_ and *Topt* in the ancestors of oviparous and viviparous species a two-step approach was used. First, the ancestral states of reproductive mode were estimated for each node in each of the 1000 trees using HMMs (R code model M5_corHMM). The phylogenetic distribution of the evolutionary origins of viviparity in squamates remains highly debated^22,41^. Our aim here was not to try to resolve this controversy, but past literature has highlighted that the rate of evolution of viviparity varies across squamates and this has important effects on ancestral reconstructions^42–44^. Not accounting for such rate variation has resulted in the ancestor of squamates being predicted to be viviparous and many reversals of viviparity to oviparity, both of which are thought to be unlikely^43,45,46^.

We therefore used HMMs, implemented in the R package ‘corHMM’ that can estimate variation in the rate of evolution of binary characters across phylogenies^47^. To do this, a number of different rate categories from one state (e.g., oviparity) to another state (e.g., viviparity) are pre-defined and then estimated across the phylogeny. The most likely number of rate categories can be identified by comparing AIC values across models with different numbers of pre-defined rate categories.

We found that on the trimmed phylogeny (224 species) AIC values were lowest when there were 2 rate categories (See R script ‘4. PBT models.R section 5’). This indicated that in two clades, transitions to viviparity occurred at a higher rate than in other parts of the phylogeny (Fig. S5). This model produced ancestral estimates that are consistent with the predominant view of viviparity evolution across squamates^43,48^: a root state of oviparity and relatively few reversals of viviparity to oviparity compared to the transitions from oviparity to viviparity (Table S9). Estimates of ancestral states from HMMs were used to identify transitions between oviparity and viviparity by classifying nodes in the following way: 1) oviparous with only oviparous descendants (oviparous to oviparous); 2) viviparous with only viviparous descendants (viviparous to viviparous); 3) oviparous with at least one viviparous descendant (oviparous to viviparous); and 4) viviparous with at least one oviparous descendant (viviparous to oviparous). In the second step, a MR-BPPM with Hedges’ *g, P*_*bt*_, and *T*_*opt*_ as response variables was used to reconstruct ancestral states for each trait (R code model M5.1). From this model mismatches in the thermal optima of females and embryos (CI of *P*_*bt*_ *– T*_*opt*_ not overlapping 0) and values of Hedges’ *g* were estimated for each node (Table S8 & S9). To test if female plasticity differed between the ancestors of oviparous and viviparous lineages, with and without mismatched mother-offspring thermal optima, we examined if estimates of Hedges’ *g* were different (CIs not overlapping 0) between matched and unmatched thermal optima for transitions from ‘oviparous to viviparous’ compared to transitions from ‘oviparous to oviparous’.

To verify that our ancestral estimates of Hedges’ *g, P*_*bt*_, and *T*_*opt*_ from the MR-BPMM were robust to variation in rates of evolution across the phylogeny we used phylogenetic ridge regression (PRR) implemented in the R package ‘RRphylo’^49^. We found that PRR models that allowed for rate variation produced similar estimates to BPMMs for each trait (Pearson’s correlation coefficient (*r):* Hedges’ *g* = 0.82, *P*_*bt*_ = 0.94, *T*_*opt*_ = 0.98. R script ‘4. PBT models.R section 5’). Given rate shifts had minimal impact on estimates of ancestral states, we used estimates from the MR-BPMMs because they: 1) allowed missing data; 2) incorporated sampling variances associated with response variables; 3) enabled phylogenetic correlations to be estimated; and 4) produced distributions of estimates (posterior samples) for each node that allowed significant thermal mismatches between embryos and adults to be calculated.

To account for phylogenetic uncertainty in estimates of viviparity from HMMs and Hedges’ *g, P*_*bt*_ and *T*_*opt*_ from MR-BPMMs we estimated the ancestral states for each trait for each node in each of the 1000 trees (Table S9). Quantifying discrepancies between embryo and adult thermal optima in relation to transitions in reproductive mode requires summarising posterior distributions of estimates of *P*_*bt*_ and *T*_*opt*_ for each node, and relating it to its transition category (oviparous to oviparous viviparous to viviparous, oviparous to viviparous, viviparous to viviparous). One complication is that for each node the predicted transition category can vary across trees due to differences in topology and internal tree structure. Discrepancies between *T*_*opt*_ and *P*_*bt*_ can be estimated for each transition category for each node, but this becomes problematic when some transition categories for some nodes are rare as it results in few posterior samples to estimate discrepancies. To circumvent this problem, each node was classified according to the most frequently predicted transition category and related to posterior distributions of Hedges’ *g, P*_*bt*_ and *T*_*opt*_ summarised across all trees.

### Discrepancies between embryo and adult thermal optima and the evolution of viviparity

To examine if viviparity evolves more frequently in lineages where the thermal optima of adults and embryos are aligned, we tested if oviparous nodes with similar thermal optima (CI of *P*_*bt*_ *– T*_*opt*_ overlapping 0) produced more descendent viviparous lineages than nodes where there were mismatches in thermal optima (CI of *P*_*bt*_ *– T*_*opt*_ not overlapping 0). Differences in frequencies were tested using a chi^2^ test of the number of nodes with and without thermal mismatches for oviparous nodes with oviparous descendants versus oviparous nodes with viviparous descendants (R script ‘5. PBT Proc.R section 5B)’

### Verification analyses

### Checking for differences in Hedges’ g between laboratory and field studies

Whether laboratory and field studies differed in their estimates of Hedges’ *g* was checked using a BPMM of Hedges’ *g* with study type as a fixed effect (R code model M5.4. Table S10).

### Checking ancestral state reconstructions of viviparity were robust to missing data

We examined how well models predicted ancestral values of reproductive mode with missing tip data using HMMs in two ways. First, we compared the ancestral states of nodes predicted using all available data on reproductive mode from Pyron and Burbrink^28^ (n_species_= 7831, Table S2) to the predicted states obtained using only the trimmed tree and data (n_species_=224). Second, we examined the accuracy with which ancestral nodes could be predicted on the phylogeny of 7831 species using only reproductive mode data from the 224 species with thermal data. The predicted ancestral states from both these analyses can be found in Table S11.

## Notes

https://doi.org/10.17605/OSF.IO/JT28V

### Competing Interest Statement

The authors have declared no competing interest.

### Summary of Updates

Text and figures revised, results unchanged

