## Supplementary material for "Maternal behavioural thermoregulation facilitated evolutionary transitions from egg laying to live birth": Fig. S

**Figures**

Records identified through literature search

N = 721

Identification

Records after duplicates removed

N = 698

Screening

Records excluded

N = 574

Screened records

(title + abstract + keywords)

N = 124

Eligibility

Full-text articles excluded, with reasons (irrelevant)

N = 47

Full-text articles

N = 77

Full-text articles excluded, with reasons (no reported sample size, error)

N = 4

Studies included in comparative analysis

N = 73

Identification

**Fig. S1**: PRISMA statement for Dataset 1- Preferred body temperature in gravid and non-gravid females and effect size calculation. Our original literature search using ISI *Web of Science* yielded 721 papers. Of these 23 were duplicates and were removed, 574 records based on the abstract and 47 full-text articles were rejected based on irrelevance – these studies did not measure both gravid and non-gravid body temperatures in female squamates. A further 4 records did not report sample size or a statistic from which sample size could be calculated and/or a measure of error from which standard deviation could be determined. A resulting 77 articles representing 54 species were included in the final analysis.

Records identified through literature search

N = 1075

Identification

Records after duplicates removed

N = 991

Screening

Records excluded

N = 627

Screened records

(title + abstract + keywords)

N = 364

Full-text articles excluded, with reasons (no mention of sex of measured individuals)

N = 214

Eligibility

Full-text articles

N = 150

Full-text articles excluded, with reasons (measured during reproductive season)

N = 8

Studies included in comparative analysis

N = 142

Identification

**Fig. S2**: PRISMA statement for Dataset 2 - Preferred body temperature in non-gravid females. Our original literature search using ISI *Web of Science* yielded 1075 papers. Of these 84 were either duplicates, or already included in dataset 1 (see above) and were removed, 627 records based on the abstract and 214 full-text articles were rejected based on irrelevance – these studies did not specify sex of the measured individuals. A further 8 records were excluded since individuals were measured during the reproductive season. A resulting 142 articles (supplemented with 42 studies from ^10^) were included in the final analysis.

Records identified through literature search

N = 402

Identification

Records after duplicates removed

N = 340

Screening

Records excluded

N = 263

Screened records

(title + abstract + keywords)

N = 77

Full-text articles excluded, with reasons (not hatching success)

N = 43

Eligibility

Full-text articles

N = 34

Full-text articles excluded, with reasons (less than 3 temperatures measured)

N = 24

Studies included in comparative analysis

N = 10

Identification

**Fig. S3**: PRISMA statement for Dataset 3 - Calculation of embryo thermal performance. Our original literature search using ISI *Web of Science* yielded 402 papers. Of these 62 were duplicates or already included from the Reptile Development Database ^4^ and were removed, 263 records based on the abstract and 43 full-text articles were rejected based on irrelevance, for example they measured another metric of thermal performance other than hatching success. A further 24 records measured hatching success across less than three temperatures, such that thermal performance curves could not be estimated. A resulting 10 articles were included in the final analysis in addition to those obtained from the Reptile Development Database.

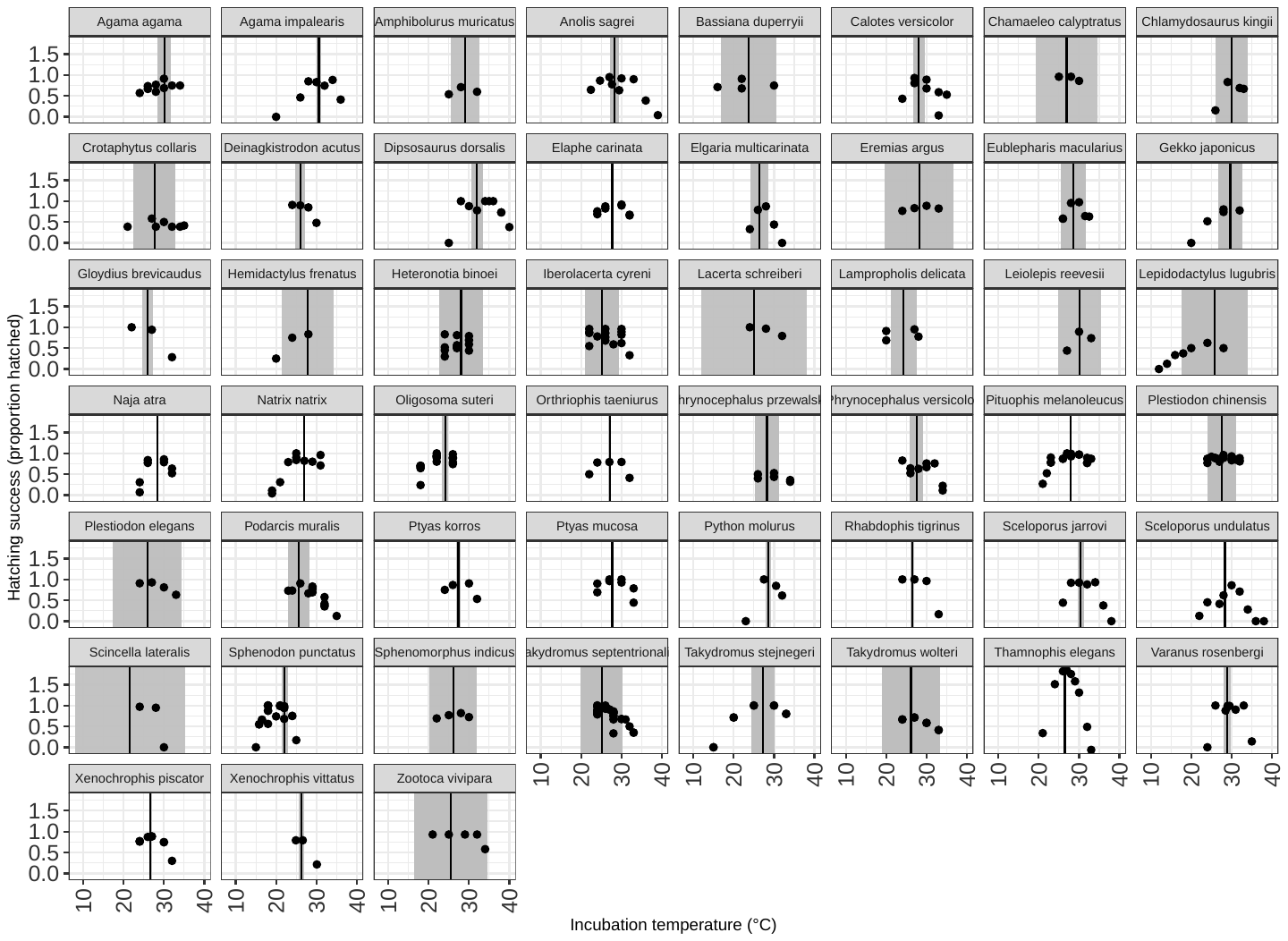

Fig. S4. Raw data used to calculate the temperature that optimizes hatching success (*T_opt_*) for 51 oviparous (n = 47) and viviparous (n = 4) species. Plots show species-level hatching success (%) across constant incubation temperatures (°C). Imputed values of *T_opt_* and sampling error from a Bayesian Phylogenetic Mixed Effects Model shown by black and coloured lines, respectively.

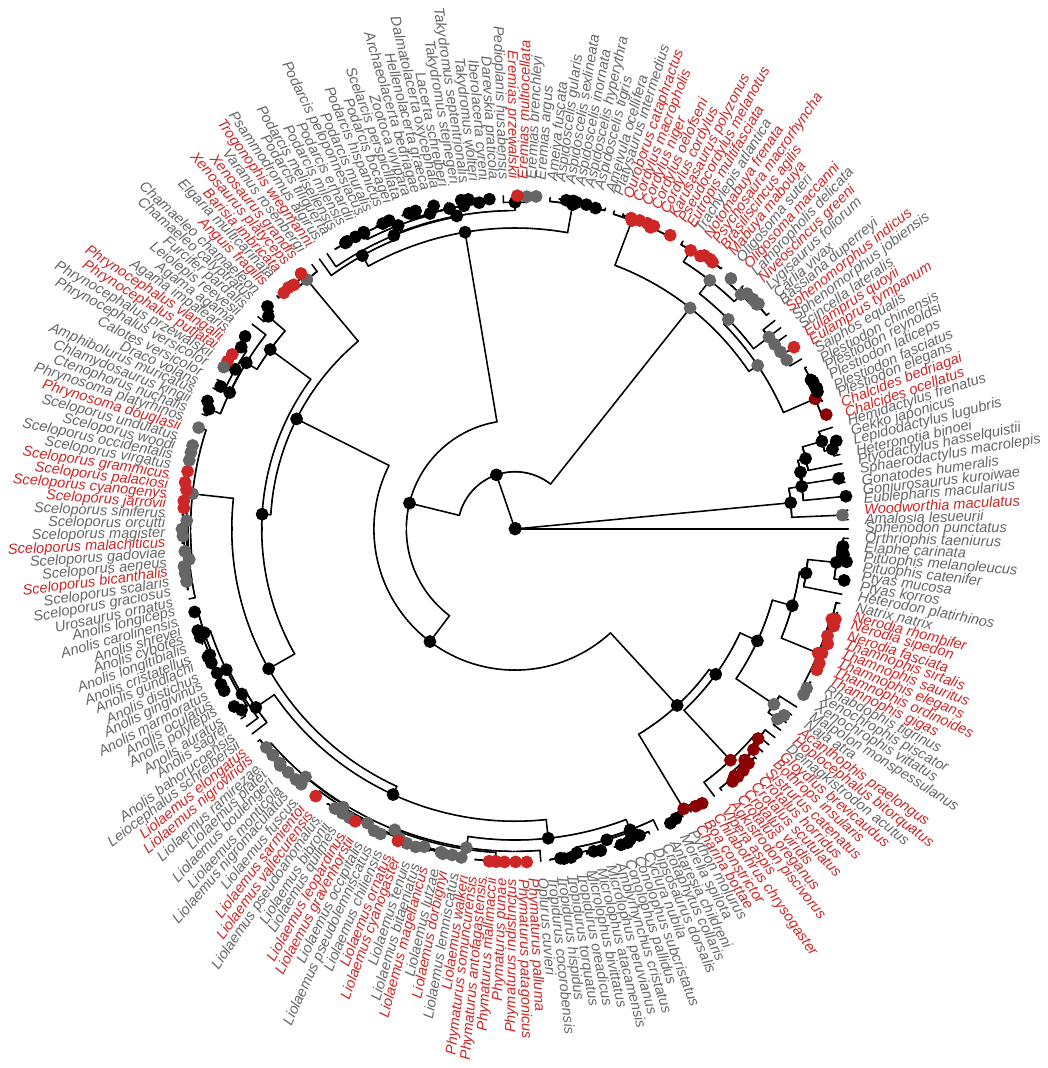

Fig. S5. Changes in the rate of evolution in viviparity across the trimmed phylogeny. Rate shifts were identified using hidden markov models using the R package ‘corHMM. Black and dark grey circles indicate oviparous states with different rates of evolution, and dark red and red circles indicate viviparous states with different rates of evolution.

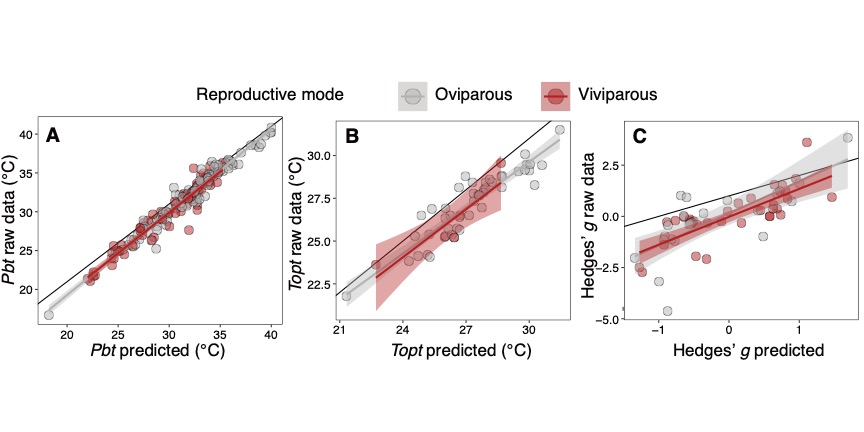

Fig. S6. The relationship between raw data and values predicted by MR-BPMMs. A) preferred body temperature (*P_bt_*), B) optimal temperature for embryos (*T_opt_*) and C) changes in female body temperature when gravid (Hedges’ *g*). Black dotted line represents a 1:1 relationship between raw and predicted values.
